# Packed hydrogel microfibers as a granular medium

**DOI:** 10.1101/2023.06.19.545582

**Authors:** Gregory S. Grewal, Georgia T. Helein, Jenna L. Sumey, Steven R. Caliari, Christopher B. Highley

## Abstract

Particle-based (granular) hydrogels are an attractive class of biomaterials due to their unique properties and array of applications in the biomedical space – serving as platforms for extrusion printing and injecting as well as permissive materials for 3D cell culture. Physical properties of particle-based hydrogels are governed in part by contact forces between particles, which are limited to interactions with neighboring particles. Secondary annealing mechanisms are often used to increase mechanical properties and serve to link particles across the granular material volume. Here, we present a novel particle-based hydrogel where each “particle” is a discrete electrospun hydrogel microfiber that has been segmented to a length of 93 ± 51 𝜇m, with a diameter of 1.6 ± 0.3 𝜇m. The fibers are flexible and have aspect ratios that are greater than one order of magnitude larger than most traditional hydrogel microparticles. This enables long-range entanglements of discrete fibers following packing into a bulk material, yielding unique properties. Without crosslinking, these packed hydrogel microfiber materials are mechanically robust, they can stretch without breaking when strained, and they exhibit stress relaxation under constant strain. As a cell culture scaffold, shear-induced alignment of the individual fibers within 3D printed filaments confers contact guidance cues to cells and promotes anisotropic cellular morphologies. Packed hydrogel microfibers can also be used as 3D cell culture environments, with cells able to spread due to the permissive nature of the scaffold. Overall, this work introduces a particle-based material system comprised of individual hydrogel microfibers that allows unique properties to be engineered into biomaterials that might be used in extrusion processes and cell cultures and, ultimately, in tissue engineering and regenerative medicine applications.

## Introduction

In recent years, granular hydrogels – which are hydrogel materials comprised largely of discrete hydrogel microparticles (HMPs) held in place by particle-particle contact forces and, often, engineered interparticle interactions – have received increasing attention in biomaterials research^1^. Granular hydrogels are attractive for reasons that include properties that enable injection delivery as well as control over mechanics and degree of porosity^2–4^. Discrete HMPs, often spherical particles formed through microfluidic or emulsion approaches (on the order of 10^1^-10^2^ 𝜇m in diameter)^1, 5^, are packed to yield a macroscale construct held together by physical (e.g., contact)^6^ or chemical (e.g., covalent)^7^ interactions, or both. Physical interactions are commonly introduced through packing of individual HMPs via centrifugation or vacuum filtration where interstitial fluid between the particles is largely removed^1, 5^. This places HMPs in direct physical contact where particle-particle contact interactions determine mechanical properties of the granular hydrogel as a whole^8, 9^. These purely physical interparticle forces allow for the granular hydrogel to be stable at rest, but individual particles will begin to slide and flow when a force is applied that overwhelms contact interactions in the system (e.g., during extrusion)^6, 10–12^.

Chemical crosslinking, or annealing, between particles can improve mechanical stability in granular hydrogel systems^13–15^. Annealing can be employ covalent^2^ or supramolecular^16^ crosslinks between reactive moieties on the surface of discrete particles to stabilize the granular hydrogel. In injection applications into a tissue defect or wound, this stabilization has enabled a class of granular hydrogels known as microporous annealed particle (MAP) scaffolds to serve as injectable tissue regeneration platforms. There has been considerable work tuning HMP properties, and therefore granular hydrogel behaviors, by varying crosslinking and secondary annealing mechanisms, introducing degradability, and using HMPs to deliver bioactive molecules post-delivery^7, 12–18^. This degree of specificity over individual particles allows for engineered modularity^19^ and control over physical and chemical heterogeneity in granular hydrogel materials.

Advanced strategies to tune the biomimicry of individual HMPs are largely predicated on spherical particles (aspect ratio ∼ 1). Shifting away from spheres, studies leveraging particles with increased aspect ratios have elucidated some unique characteristics when assembled together into granular hydrogel scaffolds. For example, rod-shaped HMPs (aspect ratios ranging from ∼ 2-20) enable larger, more interconnected pores throughout the scaffold, which facilitate greater cell migration and infiltration^20, 21^. Increasing the size of HMPs while conserving the higher aspect ratios, leads to long, flexible hydrogel strands that can align and entangle when assembled into a granular hydrogel^22, 23^ – offering interactions at increased length scales compared to other, lower aspect ratio HMPs. These entanglements are useful for extrusion mechanisms^23–25^ (e.g., injection and 3D printing) and also enable increased granular hydrogel structural fidelity without secondary annealing^23, 26^.

Decreasing the diameter of these high aspect ratio strands to sub-cellular length scales (∼1 𝜇m diameter “fibers”) offers a granular hydrogel platform where cells are able to recruit individual fibers and reorganize the structure of the scaffold^27^. While these have been densely assembled and immobilized through designed annealing interactions^27–29^, we sought to develop a materials strategy using dense combinations of hydrogel fibers without secondary annealing mechanisms to yield granular hydrogels with high mechanical stability and high degrees of permissivity. To achieve this, we electrospin and segment hydrogel microfibers to yield discrete “particles” with sub-cellular length scale diameters and high aspect ratios, which are analogous to extracellular matrix (ECM) fibrous proteins like collagen^30–32^. These individual microfiber segments represent the “grains” that are packed through centrifugation to form the granular hydrogel scaffold – hereafter denoted as packed hydrogel microfiber (PHM) scaffolds. Herein, we demonstrate that PHM scaffolds allow the design of materials which are strain yielding and while exhibiting unique stretching properties. Furthermore, we show that these materials dissipate stress in response to applied strains similar to biological materials and maintain their mechanics with increasing interstitial fluid within the system (i.e., decreasing packing density). Finally, we show that PHM scaffolds can influence cell behaviors through topographical cues, which can be dictated via extrusion and bioprinting, as well as through their unique physical properties as 3D, permissive cell culture scaffolds.

## Results and Discussion

### Preparing packed hydrogel microfiber scaffolds

Packed hydrogel microfibers were fabricated from electrospun methacrylated hyaluronic acid (MeHA), which has demonstrated biocompatibility^33, 34^ and been previously used in electrospinning^35, 36^ (^1^H NMR spectrum confirming MeHA synthesis shown in Figure S1). Electrospun MeHA fibers were designed to model endogenous ECMs both through the use of a material based on the native glycosaminoglycan, hyaluronic acid (HA), and its subsequent processing into hydrogel microfibers that mimic the natural protein fibers in the ECM^37–40^. Methacrylation enabled photomediated crosslinking to stabilize the fibers prior to hydration (Figure 1A) and, through reactive methacrylates that remain after photocrosslinking, coupling of the fibronectin-mimetic Arg-Gly-Asp (RGD) adhesive motif through a Michael addition reaction to a cysteine residue incorporated into the RGD-containing peptide^41^.

**Figure 1.**
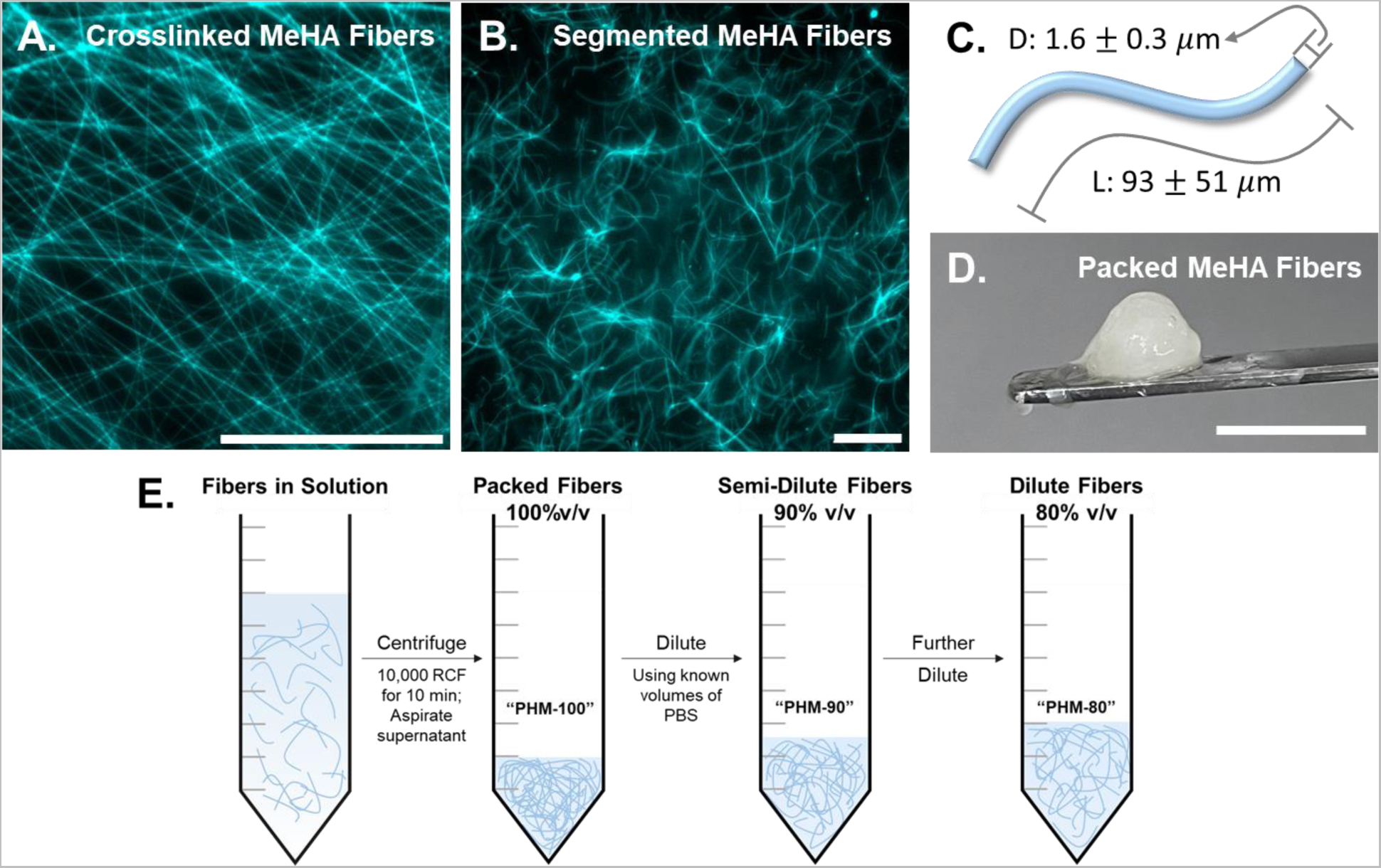
Fabrication of packed hydrogel microfiber-based hydrogel scaffolds. (A) Dry MeHA fibers containing a fluorophore for visualization immediately following the electrospinning and crosslinking processes. (B) Trituration yields fiber segments in solution that are (C) on the order of ∼100 µm in length, with diameters on the order of ∼1 µm. (D) Packing via centrifugation at 10,000 RCF for 10 minutes yields a packed hydrogel microfiber scaffold that behaves as a bulk solid at rest. (E) Schematic illustrating how PHM-100, PHM-90, and PHM-80 scaffolds were assembled, where more PBS added increases the inter-fiber fluid content while conserving total hydrogel microfiber content in each sample. *Scalebars (A-B) = 100* 𝜇*m and (C) = 1 cm*.

To create segmented microfibers that can be packed together to form PHMs, crosslinked MeHA fibers were hydrated then segmented via a series of triturations through needles of decreasing inner diameter (16 G, 18 G, and 20 G)^27, 28, 42^. The resultant fiber segments (Figure 1B) were 1.6 ± 0.3 𝜇m in diameter and 93 ± 51 𝜇m in length (Figures 1C and S2). After packing the suspension of discrete fibers by centrifugation, they behaved as a bulk solid at rest (Figure 1D), similar to conventional granular hydrogels. Compared to granular materials based on spherical particles, the high length:diameter aspect ratio of the microfibers in a PHM scaffold allows for unique long-range entanglements. These long-range interactions allow for dilution of the “fully packed” scaffolds to increase inter-fiber fluid content and provide more space between individual fiber segments. Here, we define the “fully packed” scaffolds as consisting of 100% v/v of the material recovered from centrifugation. The fully packed material, hereafter referred to as “PHM-100”, can be further diluted with known volumes of PBS to 90% v/v and 80% v/v, referred to as “PHM-90” and “PHM-80”, respectively (shown in Figure 1E).

### Mechanical characterization of PHMs

PHM-100, PHM-90, and PHM-80 all exhibit characteristics that are important for particle-based hydrogels used for extrusion processes or in cell culture systems. Shear-thinning and self-healing were evidenced in all dilutions via oscillatory shear rheology (Figure 2A) through repeated cycling between low (1%) and high (250%) strains. In contrast to granular hydrogels based on spherical particles^6, 43^, the storage modulus recovered after removal of the high strain from the system was reduced (∼60-70%) in all groups (Figure 2B). This reduction was statistically significant and observed between the first and second low-strain regimes, with no statistical significance in the percentage drop in modulus across dilutions. We attribute this PHM behavior to rearrangements of microfibers during rheometric analysis. After initial packing via centrifugation, the organization of the segmented fibers is likely maximumly entangled through random organization, giving rise to the initial mechanical properties of the scaffold. During the high-strain regimes, this organization is disrupted, and a portion of the long-range entanglements is irreversibly lost.

**Figure 2.**
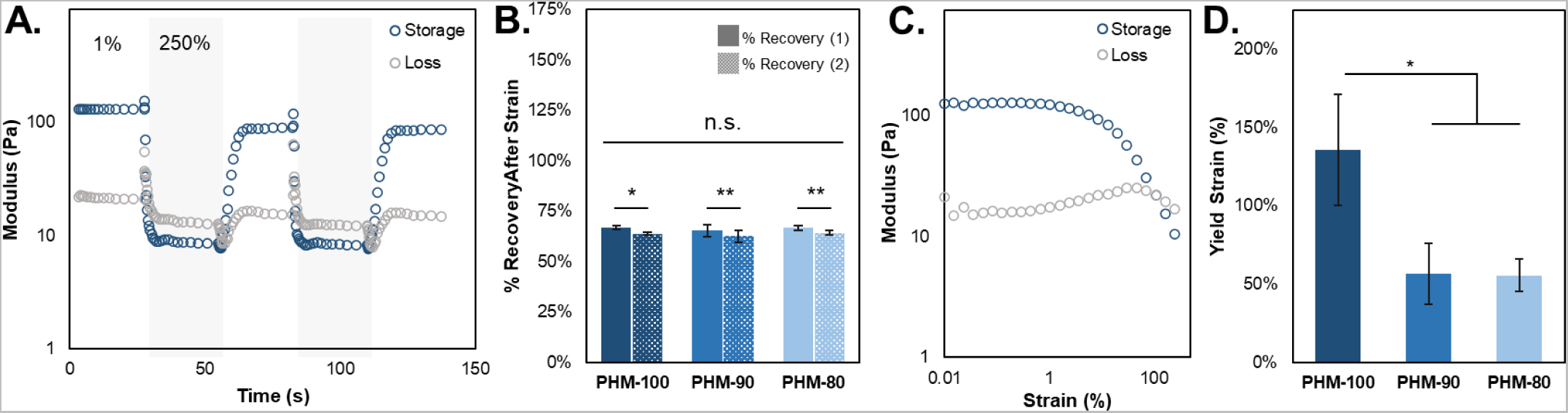
Rheological characterization and macroscale extrusion of microfiber particle-based hydrogels. (A) Cyclical application of high (250%, shaded regions) and low (1%) strains demonstrates shear-thinning and self-healing properties of PHM-100 (PHM-90 and PHM-80 shown in Figure S3). (B) Quantification of recovery after strain shows that PHM scaffolds recover ∼60-70% of their initial modulus prior to the addition of high strain – suggesting that organization of discrete fibers influences overall mechanical properties. (C) Amplitude sweep for PHM-100 demonstrates shear-yielding properties of this group with (D) yield strains for diluted groups (PHM-90 and PHM-80 rheological data shown in Figure S3) significantly less than PHM-100. *\* p < 0.05; ** p < 0.01*.

As in other granular hydrogel and shear thinning systems, the ability to convert from a solid to liquid-like state above a yield strain (defined as the crossover point between G’ and G” in a strain sweep^17^, Figure 2C) allows the solid-like material to be injected or extruded. The 136% yield strain (Figure 2D) for the 100% v/v group (PHM-100) is notably larger than other granular hydrogel systems^20^. Diluted fiber density in the PHM-90 and PHM-80 groups yielded statistically decreased yield strains of 56% and 57%, respectively, which is consistent with yield strains of other reported granular hydrogel systems^20^. These data indicate that all test groups are shear-thinning and self-healing, with yield strains generally in ranges characteristic of granular hydrogels used in extrusion processes. Furthermore, increased yield strains are possible in PHM formulations, which indicate the potential for enhancing the stability of particle-based hydrogels through dense fiber-based formulations.

### Evaluating the extensibility of PHM scaffolds

Anticipating that long-range interactions between microfibers in PHM scaffolds resulting from their high aspect ratios would offer unique bulk properties, we next examined the extensibility of the PHM-based materials. We conducted a modified version of filament stretching extensional rheology (FiSER)^44^, in which a material can be stretched as a filament, allowing strain to failure to be observed as a measure of material extensibility or stretchability. We hypothesized that discrete fibers within the filament would participate in interactions at extended length scales compared to conventional granular materials, thus resulting in highly extensible PHM materials when stretched. Compared to spherical particle interactions, which would be restricted to engagements with a limited number of neighboring particles^1^, we expected that a PHM scaffold would stretch more and appear less brittle as a bulk than a conventional granular hydrogel.

In modified FiSER testing, all groups of PHMs exhibited strain-to-break values of 2000-2500% (Figure 3A), indicating fibers maintain filament-stabilizing interactions in response to stretching. Notably, while PHM-100 exhibited the highest degree of stretchability (∼2500%), there was no statistical difference between the groups. In these measurements, the normal forces sustained by the filaments during extension (Figure 3B) were observed to decrease with respect to hydrogel microfiber density. These observations combine to suggest that the extensibility of a PHM scaffold is dictated by fiber geometry, while the density of fiber-fiber interactions (i.e., the combination of entanglements and surface-surface interactions) drives the forces that can be sustained during stretching. Correspondingly, the normal force profiles for each group exhibit the same trend as they are stretched to failure, which occurs at % strains that exhibit no statistical difference. We believe this extensibility (visualized during testing of PHM-100 in Figure 3C) is unique among particle-based hydrogels where there is no dissolved polymer between particles, and that it is driven by enhanced interactions among the discrete elements of the bulk material that result from the high-aspect ratios of the individual fibers in the PHM scaffold.

**Figure 3.**
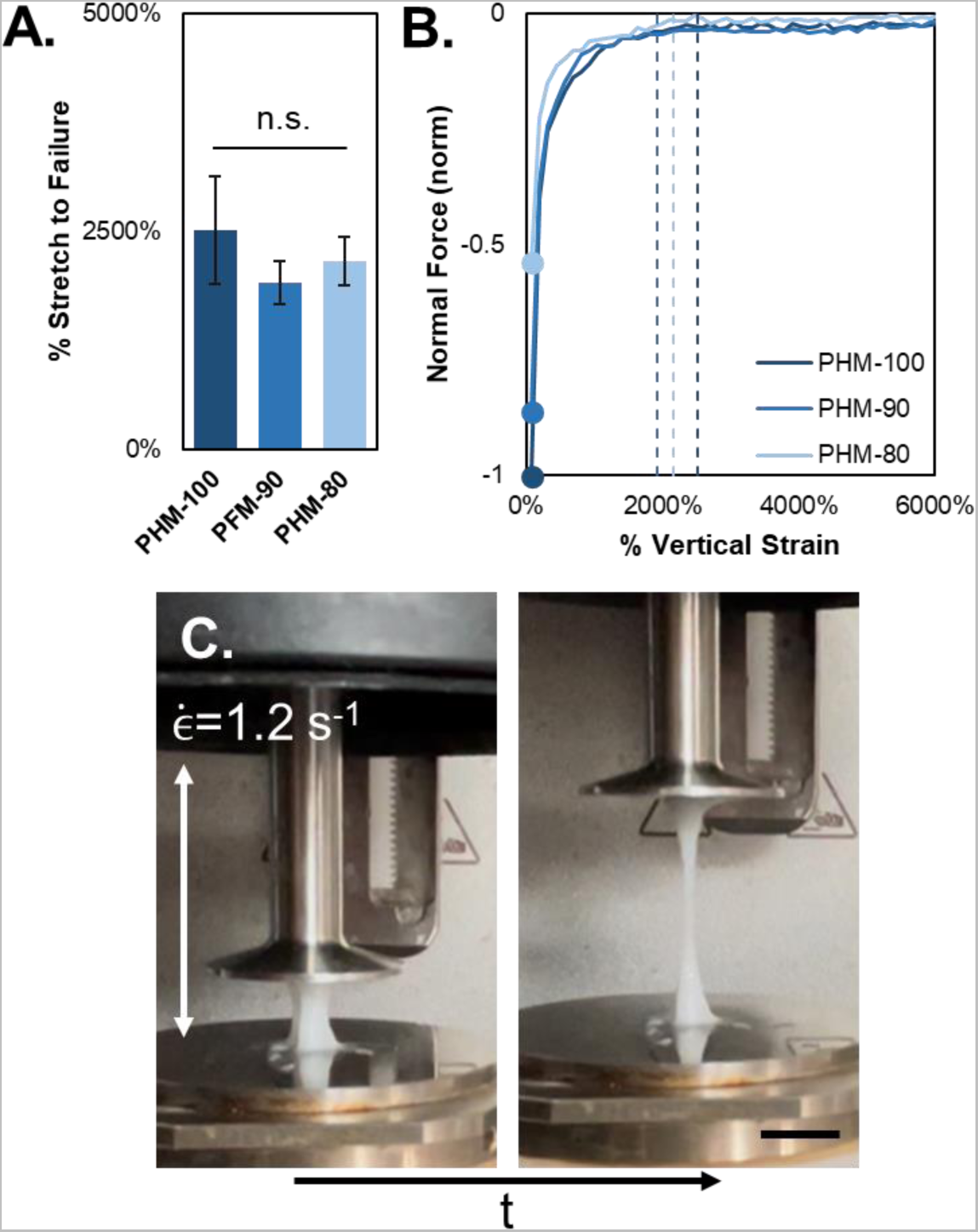
Modified FiSER characterizes extensibility of packed hydrogel microfiber scaffolds. (A) Percentage stretch to failure of PHM-100, PHM-90, and PHM-80 filaments indicates that all groups can stretch vertically to greater than 2000% of their original height with no statistical significance between groups. (B) Representative trends of normal force for each dilution (normalized to PHM-100) illustrates that although percentage stretch to failure is similar for all groups, the original normal force (indicated by filled circles at 0%) decreases as dilution increases. However, normal forces exhibit similar trends once the stretching begins. Dashed lines correspond to the average % stretch to failure values from A. (C) Representative images of PHM-100 being stretched using the modified FiSER setup. *Scalebar (C) = 1 cm*.

### Characterizing viscoelasticity and stress relaxation of PHM scaffolds

Given observation of these dynamic behaviors of PHM hydrogels, we were interested in further characterizing their viscoelasticity and stress relaxation to determine whether they might offer new properties as cell and tissue culture scaffolds. Endogenous tissue exhibits a host of complex, time-dependent mechanical properties that are difficult to recapitulate in traditional hydrogel materials^45–47^. For example, ECM structural components like collagen fibers present in many native tissue microenvironments enable the dissipation of stress over time, thereby contributing to nonlinear viscoelasticity profiles^48^. Inspired by this and other long-standing observations of the similarities between the geometries of electrospun fibers and ECM fibrous proteins^30^, we hypothesized that noncovalent interactions between individual fibers in 3D PHM materials would allow effective mimicking of the stress relaxation of native ECM in a synthetic system. As noted above, the previous characterization experiments suggested the ability of fibers within the PHM hydrogels to interact at rest but slide past each other and reorganize to dissipate forces applied to the materials.

To assess PHM scaffold viscoelasticity for comparison to biological tissues and materials^45^, we measured the loss modulus (i.e., viscous component, G”) versus storage modulus (i.e., elastic component, G’) from rheometric time sweeps (Figure 4A). All PHM formulations exhibited relatively soft bulk storage moduli – ranging from ∼50 Pa for PHM-80 to ∼150 Pa for PHM-100, despite individual hydrogel microfibers having moduli many orders of magnitude greater than bulk fiber-based scaffolds^36^. Additionally, all groups demonstrated viscoelastic behavior with their storage moduli approximately 5x their loss moduli (grey dashed line in Figure 4A illustrates where 𝐺^′^ = 5 × 𝐺′′). Importantly, endogenous tissue typically possesses a higher elastic contribution, where the storage moduli are typically 10x the loss moduli^45^. We further postulated that we could achieve this 10x target via additional interfiber crosslinking (or annealing) of PHMs.

**Figure 4.**
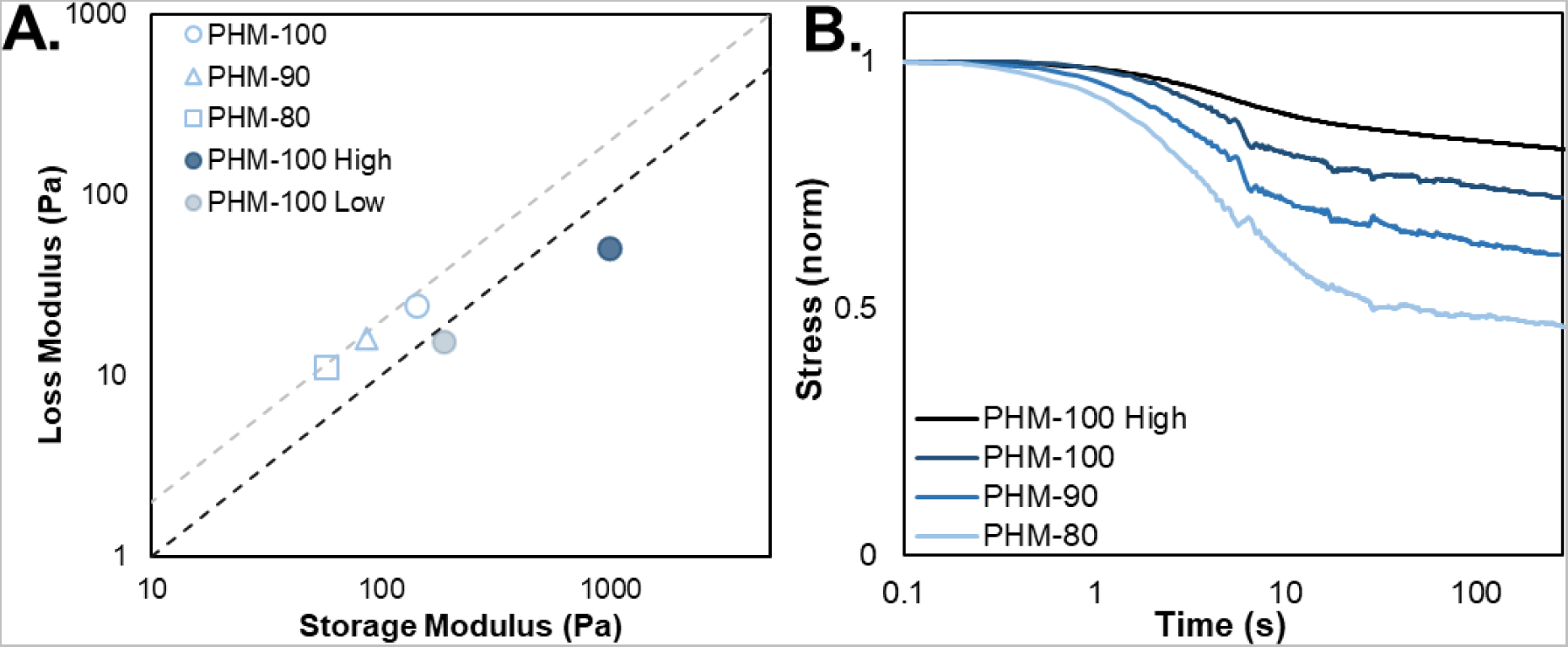
Characterization of viscoelasticity and stress relaxation properties. (A) Plot of shear loss modulus versus shear storage modulus of PHMs. Grey dashed line and black dashed line represent where G’ is equivalent to 5x G’’ and 10x G”, respectively. Non-crosslinked groups reside near the 5x trend, whereas the addition of secondary annealing to stabilize the material shifts the viscoelasticity to the 10x trend, which is characteristic of many biological tissues^45^. (B) Plot of stress relaxation tests where a 15% constant shear strain was applied to the systems and resultant stress was observed as a function of time. All groups exhibit relatively rapid stress relaxation and mimic profiles characteristic of viscoelastic solids. PHM-100, PHM-90, and PHM-80 are compared with the PHM-100 High group.

To test this, we designed PHMs to have residual methacrylate groups on the surfaces of individual microfibers that were not consumed during the initial fiber crosslinking process. Utilizing the PHM-100 formulation, we created “High” and “Low” crosslinkable fibers, where the “PHM-100 High” group was treated after fiber segmentation to remove some reactive methacrylate groups (to mimic RGD modification) but had most residual methacrylate groups available for crosslinking. The “PHM-100 Low” group was similarly treated after fiber segmentation to eliminate most, but not all, residual methacrylate groups on the fibers. Therefore, PHM-100 High could achieve considerable annealing during a secondary crosslinking step by inducing methacrylate polymerization. Conversely, PHM-100 Low could undergo only a comparatively low degree of secondary crosslinking, resulting in reduced interfiber annealing compared to PHM-100 High (rheology for the secondary UV-initiated annealing step shown in Figure S4).

We observed in both groups (PHM-100 High and PHM-100 Low) that interfiber crosslinking shifted the G’:G’’ ratio towards 10:1 (Figure 4A, shaded circles, dashed black line represents 𝐺^′^ = 10 × 𝐺′′). PHM-100 Low exhibited a stark decrease in G” coupled with a marginal increase in G’ compared to PHM-100. Interestingly, PHM-100 High exhibits a slight increase in the loss modulus coupled with a notable increase in the storage modulus, which also shifted the viscoelastic moduli toward the 10:1 ratio that characterizes many biological tissues. Taken together, these data suggest that the degree of secondary crosslinking afforded by available methacrylate groups can be utilized to controllably modulate the viscoelasticity of PHM hydrogel systems.

As previous tests pointed to dynamic responses to applied stress in PHM scaffolds, we next sought to evaluate their capacity to undergo stress relaxation. Time-dependent stress relaxation is a feature of many biological material systems that is observed to critically influence cellular behaviors^45, 49^; however, it is challenging to engineer and control stress relaxation in many hydrogel systems. In this analysis, we applied a constant shear strain of 15% to the sample and recorded the resultant stress as a function of time to assess how the material relaxes in response to the applied strain. Plotted as the normalized stress in Figure 4B, PHM-100, PHM-90, and PHM-80 all exhibit varying degrees of stress relaxation, with increasing relaxation corresponding to increased dilution with PBS. The enhanced relaxation for the diluted groups is attributed to more space for fibers to reorganize in response to the applied strain and reduced interactions between fibers on a per volume basis. PHM-100 experiences the highest degree of these interactions and the most confining interfiber space that slows or prevents fiber movement that results in PHM stress relaxation.

To observe the effects of interfiber crosslinking, we applied the same 15% strain to the PHM-100 High group. We observed that PHM-100 High dissipated stress modestly (Figure 4B). However, the covalent interfiber crosslinking restricted the ability of the microfibers to move and thus prevented the material from relaxing to the extent of the non-crosslinked groups (Figure 4B). Importantly, with respect to recapitulating the relaxation time scales of viscoelastic solids, the relaxation profiles of the PHM hydrogels began to plateau within 100 s of stress being applied. Through the use of a viscoelastic standard linear solid model (Figure S5), characteristic relaxation times (𝝉) were calculated^50^. All groups, including the PHM-100 High crosslinked material, exhibited most of their stress relaxation within a characteristic time of approximately 5-10 s, indicating that PHM systems respond to applied strains rapidly. From these data, we expect these non-crosslinked materials to be useful in engineering soft tissue systems *in vitro*, and in engineering both soft and stiffer tissues (via secondary crosslinking) to model physiological systems where ∼100s and quicker relaxation times are desired.

### Extrusion of PHM inks

As mentioned, the rheological and mechanical properties of PHM materials should enable diverse uses in applications where injectable or extrudable biomaterials are desired, including in biomedical applications of 3D printing. We studied the responses to extrusion processes using a 3D printer (FELIX BIOPrinter). In controlled extrusions, we first printed a 2 cm vertical filament of PHM-100 through a 22 G needle (ID: 0.413 mm). This yielded a filament with a diameter of ∼0.5 mm (Figure 5Ai). Towards observing the stability of PHM filament without interfiber crosslinking (a filament stabilized strictly by noncovalent interactions), we translated the vertical filament 1 cm horizontally without further extrusion (Figure 5Aii) and then back to the original position (Figure 5Aiii). The filament was easily manipulated without breakage. Additionally, the filament stretched noticeably as a result of undergoing dynamic stress relaxation when extended without extrusion (Figure 5Aiii, dashed circle). To further observe the extent to which the filament was extensible, the nozzle was moved vertically an additional 2 cm without extrusion. We attribute this high degree of extension to the long-range interactions among the individual microfibers, which provide additional stability in the filament that would not occur with other particle-based systems with smaller aspect ratios.

Finally, towards demonstrating the remarkable stability of the non-covalently annealed PHM filaments, we extruded filaments horizontally across the posts of a 2.5x2.5 cm inverted table (Figure 5B). The resultant filaments were mechanically stable, spanning these gaps without breaking while stabilized only by physical interparticle interactions. This demonstration of the strength of long-range interactions among individual fibers being sufficient to maintain filament integrity at longer scales is an exciting feature of the PHMs as a granular hydrogel system. These properties will have value in numerous applications that use extrusion processes, including 3D printing, where structural fidelity of printed hydrogel inks remains a central consideration in developing new biomaterial inks.

**Figure 5.**
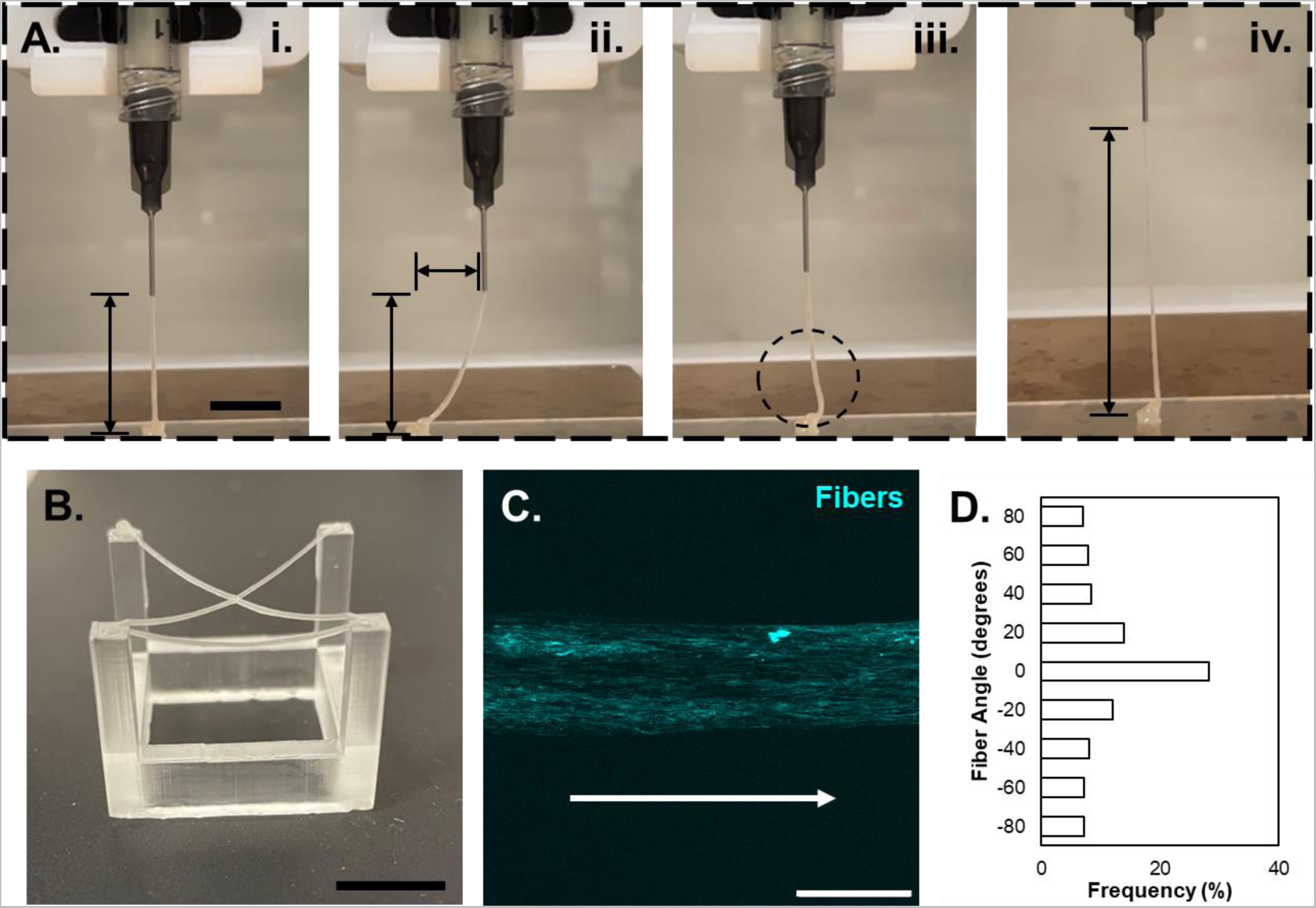
Extrusion of packed hydrogel microfiber inks. (A) (i) PHM-100 was extruded at a rate that yields a 2 cm vertical filament that is ∼0.5 mm in diameter. (ii) Following extrusion, the filament was then manipulated to demonstrate the fidelity of PHM-100. The filament was translated 1 cm horizontally before (iii) returning to the original horizontal position. The dashed circle highlights filament sagging due to stretching. (iv) The filament was finally stretched another 2 cm vertically without breaking. (B) Macroscale extrusion of PHM-100 across 4 posts of a 2.5x2.5 cm table without secondary crosslinking. These demonstrations highlight the shear-thinning and self-healing properties that are desired for extrusion printing, with the high extensibility and long-range entanglements enabling stretching and filament fidelity at long ranges without additional annealing mechanisms. (C) Fluorescent image of the microscale topography of a printed filament illustrating shear-induced alignment (white arrow indicating direction of printing) of the fibers following extrusion. (D) Quantification of fiber direction indicates that a high percentage of fibers are aligned in the direction of shear (0 degrees corresponding to the direction of the white arrow). *Scalebar (A-B) = 1 cm, (C) = 500* 𝜇*m*.

In addition to observing these macroscale characteristics of extruded filaments, we were interested in the effects of extrusion processing on the PHM filaments at the microscale, in particular the organization of the fibers after extrusion. From our previous rheological studies, we concluded that the organization of fibers within PHM scaffolds drives mechanical properties but can be influenced by high shear. Therefore, we hypothesized that the shear introduced by the extrusion process might disrupt fiber orientation and induce anisotropic alignment in the direction of shear. To assess this, we mixed nonfluorescent fibers with fluorophore-tagged fibers at a 10:1 ratio to enable visualization of the extruded material, and printed PHM-100 onto glass coverslips. Indeed, fibers aligned in the direction of the applied shear (Figure 5C-D). This result follows previously demonstrated shear-induced alignment of fibers embedded in bulk gels by Prendergast and coworkers^42^, as well as alignment demonstrated by Kessel et al. using hydrogel microstrands^23^ and Sather et al. utilizing self-assembled supramolecular nanofibers^51^. However, in the PHM materials used here, which consisted entirely of electrospun fibers with diameters on the order of 1 𝜇m, there was a unique opportunity to directly control the surface topography to direct cell behaviors. The importance of microscale topography and contact guidance on cellular behaviors has been well studied and characterized^52–55^, and the ability to dictate surface topography via extrusion using PHMs is an exciting opportunity to extend work using electrospun hydrogel fibers.

### PHM scaffolds support 2D and 3D cell culture

Given the potential to use PHM materials in bioprinting to create materials with specified surface anisotropies and to design permissive 3D environments using PHMs, we next sought to examine cellular responses to these materials. To assess whether fiber alignment in filaments would provide microscale topographical cues to cells that interacted with these materials, we printed a PHM biomaterial ink and seeded cells onto it. We extruded PHM-100 with 1 mM RGD (analogous to PHM-100 High) that could undergo a secondary interfiber crosslinking onto glass coverslips and then irradiated with light to stabilize the filament through secondary crosslinking. Immortalized murine myoblasts (C2C12s), which are known to respond to alignment cues^52^, were seeded atop the crosslinked PHM-100 filament (Figure 6A). Following a 2 d culture period, cytoskeletal staining showed that cells aligned with the direction of microfibers in the filament (Figure 6B-C), indicating that the microscale topography provided contact guidance that can influence cellular organization. Because fiber alignment responds to changes in needle direction (Figure S6), these results suggest the ability to influence cell directionality via the needle path during extrusion, with arbitrary 2D topography and cell alignment defined by the extrusion printing process when using PHMs as a biomaterial ink.

**Figure 6.**
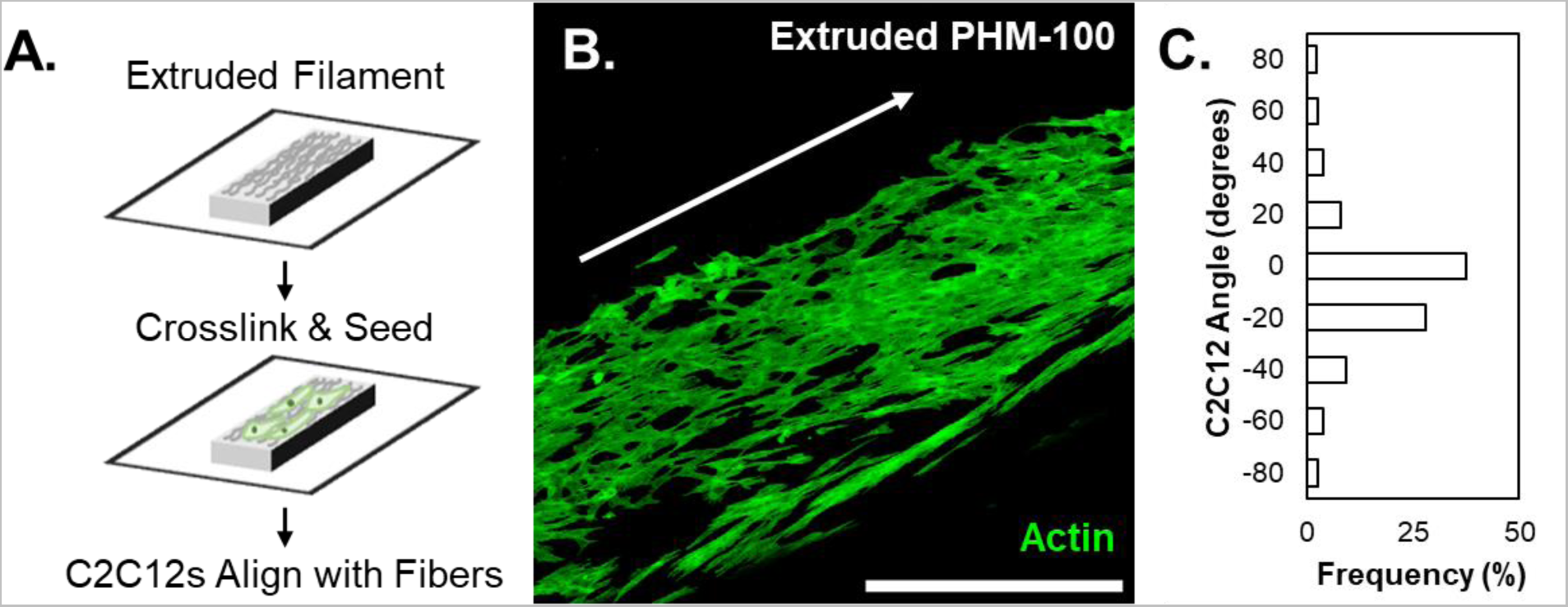
Extruded PHM scaffolds influence cell culture. (A) Process schematic for extruding PHM-100, crosslinking the individual fibers together to stabilize the filament, then seeding C2C12 cells on top of the scaffold. (B) Fluorescent micrograph of C2C12s tagged with AlexaFluor-488 phalloidin for visualization of cell alignment. (C) Quantification of cell alignment with 0 degrees corresponding to the direction of shear (white arrow). The microscale topography of the extruded filament induces alignment of C2C12 cells cultured on top of the scaffold.

We next looked at how cells within a 3D PHM-based hydrogel would behave, given the dynamic properties measured previously. Since our most fiber-dense formulation (PHM-100) is soft and viscoelastic, we postulated that C2C12s would be able to reorganize the constituent fibers during proliferation, migration, and interactions with the environment. Over short time scales, we expected to see cell spreading within these materials as opposed to rounded morphologies typical of cells in 3D hydrogels that are not permissive^56, 57^. To investigate this, C2C12s were gently mixed with PHM-100 which would undergo no interfiber crosslinking, in order to maintain the dynamic properties observed in previous experimentation. The C2C12+ PHM-100 was then placed into a PDMS mold and covered with 100 𝜇m pore filter paper to prevent PHM-100 from disassociating in culture media (Figure 7A). After 1 d in culture, cytoskeletal staining showed that C2C12s were able to spread freely in PHM-100 (Figure 7B), with a wide range of projected cell areas, along with a loss of circularity (cell shape index) that would be expected in a covalently crosslinked hydrogel quantified in Figure 7C-D.

**Figure 7.**
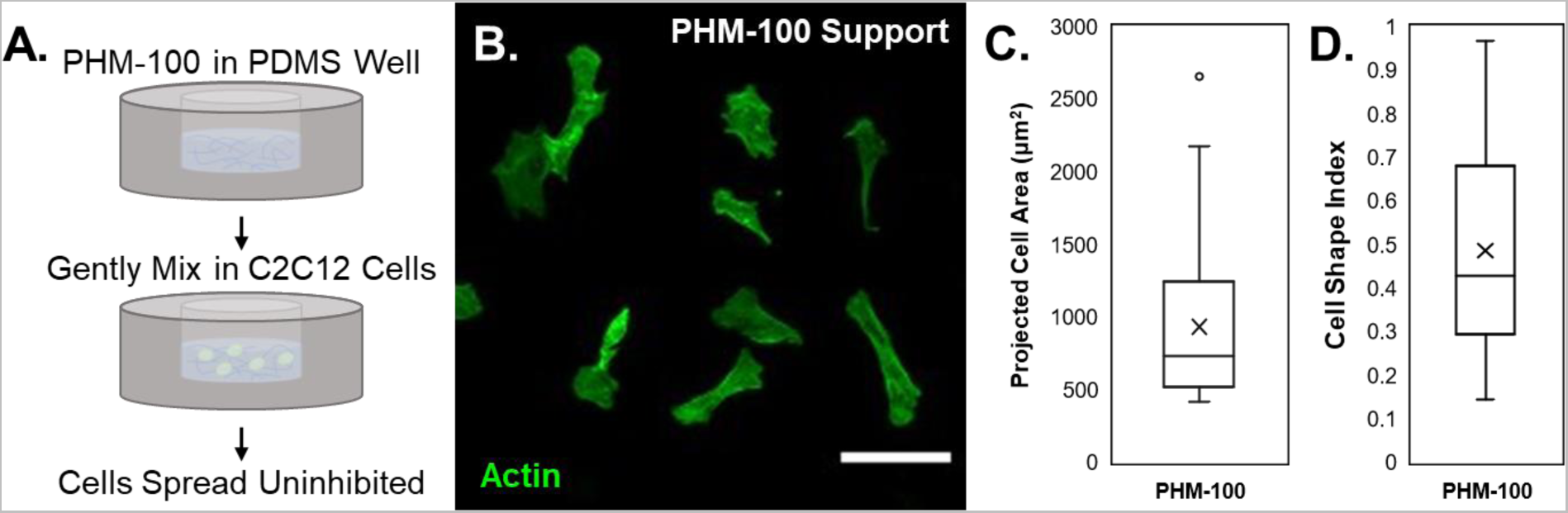
PHM scaffolds support 3D cell culture. (A) Process schematic for culturing C2C12 cells within a PHM-100 support using PDMS wells to contain the scaffolds. (B) C2C12s stained with AlexaFluor-488 phalloidin (1 d) spreading in the non-crosslinked PHM-100 scaffold. (C-D) Quantification of projected cell area and cell shape index reveal ranges of cell spreading and circularity, further supporting the permissivity of PHMs at short timescales. *Scalebar (B) = 500* 𝜇*m, (E) = 100* 𝜇*m.*

Our observations suggest that PHM-100, even absent interfiber crosslinking, supports 3D cellular activity. The cell spreading observed was in our most fiber-dense formulation, suggesting that PHMs yield in response to cells’ movements and impose minimal spatial restrictions on the cells. We believe that because the microfiber elements of the material are soft and have diameters on the order of 1 µm, they might be uniquely permissive to cellular activity. In comparison to larger spherical particles used in other granular materials, whose diameters are on the order of 10^1^-10^2^𝜇m, PHMs may present an alternative physical environment to the cells incorporated into them *in vitro* or with cells that engage with them when applied *in vivo*. We postulate that opportunities to reorganize and move throughout 3D space exist in PHM-type systems that might be less accessible in granular materials with larger particle diameters, where particle movement in response to cellular activity would be limited, resulting in cells negotiating the surfaces of the particles and the spaces in between. PHMs offer exciting new opportunities within particle-based materials through their presentation of robust bulk properties emerging from a microenvironment comprised of individual fibers that cells can readily interrogate, reorganize, and migrate around.

## Conclusions and Future Outlook

In summary, we have developed a new class of particle-based hydrogels comprised solely of discrete electrospun microfibers. Packed hydrogel microfiber scaffolds (PHMs) are shear-thinning and self-healing, with strain yielding responses that will enable application in extrusion or injection processes. The high-aspect ratios of the fibers and long-range entanglements provide physical interactions in PHMs that result in robust materials that behave elastically as a bulk below a yield strain. These interactions also enable high degrees of extensibility in packed scaffolds. PHMs can be stretched to greater than 2000% of their original height; a phenomenon that is conserved across PHM scaffolds during dilution. PHM scaffolds are also viscoelastic and quickly (∼10 s) dissipate stresses applied to them. Macroscale extrusion printing demonstrates filament fidelity and robust stability, even without secondary crosslinking to stabilize filaments. Shear-induced alignment of component microfibers during extrusion can be leveraged to direct cellular alignment when seeded on top of printed filaments. Cells seeded within these materials – or potentially infiltrating into these materials – experience permissive, 3D environments that they can interrogate and possibly remodel during migration and proliferation.

We envision this novel packed hydrogel microfiber system will provide an exciting new avenue for designing granular hydrogel media. The flexibility afforded by the PHM preparation process coupled with the ability to tailor individual fibers that comprise the bulk scaffold will enable new approaches to engineering synthetic models of the ECM *in vitro* and new opportunities for engineering implantable biomaterials.

## Experimental Section/Methods

All reagents were purchased from Millipore Sigma, unless otherwise stated.

### Methacrylated hyaluronic acid (MeHA) synthesis

Hyaluronic acid (HA) was functionalized with methacrylates as previously discussed^58^. Briefly, sodium hyaluronate (Lifecore, 60 kDa) was dissolved in deionized water at 2% w/v. While maintaining the solution at a pH of ∼ 8.5-9, methacrylic anhydride (Sigma Aldrich, 4.83 mL per g HA) was added dropwise to the solution. The reaction mixture was maintained at a pH of 8.5-9 for 6 h on ice, then continued to react at room temperature overnight. The reaction was dialyzed against deionized water (SpectraPor, 6-8 kDa molecular weight cutoff) at room temperature for 5 days, then frozen and lyophilized to dryness. The final methacrylate functionalization was 100% by quantification with ^1^H NMR (500 MHz Varian Inova 500) (Figure S1).

### Electrospinning MeHA microfibers

To electrospin MeHA, solutions consisting of 3% w/v MeHA, 2.5% w/v polyethylene oxide (900 kDa), and 0.05% w/v 2-hydroxy-4’-(2-hydroxyethoxy)-2-methylpropiophenone (HHMP) were mixed overnight in DI H_2_O. To fluorescently tag microfibers for characterization and visualization, 0.4% w/v fluorescein isothiocyanate-dextran (FITC-dextran) was included in the electrospinning solution. The MeHA solution was extruded through a 16-gauge needle at a rate of 0.5 ml h^-1^ with an applied voltage of 12-14 kV. Microfibers were collected on a negatively charged (−4 kV) rotating mandrel (DOXA Microfluidics) moving at a linear velocity of 10 m s^-1^. Fiber batches were collected for 1 h before a 2 min UV crosslinking step (365 nm, 5 mW cm^-2^, VWR UV Crosslinker) to stabilize fibers for subsequent segmentation steps.

### Preparation of PHMs

Fibers were hydrated in PBS for at least 1 h and then comminuted through a series of extrusion steps to yield small fiber segments. Beginning with a 16-gauge needle, the fiber solution was passed up and down the needle 25x to preliminarily break up the fibers. This process was repeated with an 18-gauge needle, and finally a 20-gauge needle to yield the final fiber segments. The resulting solution was then centrifuged, the supernatant was discarded to remove the PEO and unreacted HHMP, and the fibers were resuspended in a known volume of PBS to yield a stock concentration of 10% v/v. Fiber stock solutions were stored at 4 °C until further use. Fluorescently tagged MeHA fibers were diluted and imaged on a Leica DMi8 widefield fluorescence microscope to characterize fiber diameter and length (n>150 fibers) post-segmentation.

For cell culture assays, fibers were functionalized with a fibronectin-mimetic Arg-Gly-Asp (RGD)-containing adhesive peptide (GCGYG<u>RGD</u>SPG, Genscript) via the thiol-Michael addition reaction to promote integrin-mediated cellular interactions in the scaffold. Briefly, fibers were suspended at 10% v/v in the presence of 1 mM RGD, the pH of the solution was elevated to 8 using triethanolamine and allowed to react for 2 h at 37 °C. Fiber solutions were centrifuged and resuspended in PBS thrice to removed unreacted molecules prior to packing.

Fiber solutions were packed (condensed into a minimal volume with minimal solvent between the fibers) via centrifugation at 10,000 RCF for 10 min, with all remaining supernatant decanted to yield the shear-thinning and self-healing PHMs used within this work. PHMs were either used as is (denoted as PHM-100), or further diluted with known volumes of PBS to yield 90% v/v and 80% v/v (denoted as PHM-90 and PHM-80) relative to the original packing density to increase interstitial space between fibers.

### Preparation of crosslinkable PHMs

For low crosslinking, fibers were treated with 250 mM cysteamine at pH 8 for 2 h at 37 °C to quench most pendant methacrylate groups and prevent significant crosslinking. For high crosslinking, the fibers were treated with 1 mM cysteamine at pH 8 for 2 h at 37 °C to mimic the RGD addition scheme, while leaving most pendant methacrylates free. Crosslinked fiber scaffolds were prepared by suspending the fiber segments at 10% v/v in a 0.1% w/v HHMP solution, packed via centrifugation as previously described, handled for the experiment (e.g., rheology or extrusion), then treated with 2 min of UV light at 5 mW cm^-2^ to induce crosslinking of the unquenched methacrylate groups.

### Oscillatory rheological characterization

Mechanical properties of PHMs were determined via rheological measurements (Anton Paar MCR 302 rheometer) using a 25 mm parallel plate geometry, a solvent trap, and a gap height of 200 𝜇m. Shear-thinning and self-healing properties were determined using oscillatory time sweeps that cyclically changed between low strain (1%) and high strain (250%) at 1 Hz. To assess the ink/support properties of PHMs for extrusion-based printing applications, strain sweeps (0.01-250%, 1 Hz) were conducted to elucidate yield strains (%). Rheological measurements were conducted in triplicate. Shear recovery for each dilution was analyzed using a paired T-test and dilutions compared against each other were assessed using a one-way ANOVA coupled with a Tukey HSD post-hoc test to determine statistical significance.

For characterization of PHM viscoelasticity, time sweeps were utilized to quantify storage and loss moduli. To assess the ability of PHMs to stress relax, a constant 15% strain was applied to the system and the resultant shear stress was recorded over time. Stress relaxation plots were smoothed using an exponential smoothing algorithm for clean presentation. For low and high crosslinkable scaffolds, the samples were irradiated for 2 min at 5 mW cm^-2^ before conducting the rest of the rheological measurements. Time sweeps for UV crosslinking of PHMs are shown in Figure S4. Viscoelasticity rheological assessments were conducted in duplicate, with representative data shown.

### Extensional rheological characterization of PHMs

Extensibility of PHMs was determined via a modified filament stretching extensional rheology protocol^59^ (FiSER, Anton Paar MCR 302 rheometer) using a 25 mm parallel plate geometry. Briefly, 500 𝜇l of PHM scaffold was added to the rheometer stage and the gap height was lowered to 1 mm. A vertical strain rate of 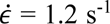 was applied, and the normal force was tracked as the geometry height was increased. Filament failure was determined at the point where the material completely broke, and the height of failure was utilized to quantify the overall percentage stretch to failure relative to the original height. FiSER experiments were conducted in triplicate, and statistical significance was evaluated using a one-way ANOVA coupled with a Tukey HSD post-hoc test.

### Extrusion printing of PHMs

A FELIX BIOprinter was used for all controlled extrusion of PHMs, and PHM-100 was utilized as the ink due to its robust mechanical properties compared to other dilutions. Inks were loaded into 1 ml syringes (BD) equipped with a 22 G needle via centrifugation at 200 RCF for 1 min to ensure all air bubbles were removed prior to printing. Printer workflows were written using G-code commands that were actuated through the FELIX BIOprinter’s software interface. Targeting filaments with 500 𝜇m diameter, macroscale properties were determined by manipulation of the filament, and microscale properties were determined by printing PHM-100 on glass coverslips. For printed filaments used in cell culture, the glass coverslips were first modified to present methacrylate groups based on a previously described protocol^60^ to allow for covalent conjugation of the filament to the coverslip. Videos of extrusion printing are included in Figures S7-S8.

### Cell culture

Immortalized murine myoblasts (C2C12s, ATCC) were used for cell culture experiments (passages 6-7). Cells were cultured in standard growth media comprised of high glucose Dulbeccos’s Modified Eagle’s Medium (DMEM), 10% v/v fetal bovine serum (Gibco), and 1% v/v antibiotic/antimycotic (Gibco). Media was changed every 2 days during expansion and experiments.

For experiments with cells seeded atop printed filaments, the crosslinked scaffolds were first sterilized via irradiation with germicidal light for 2 h. Scaffolds were then hydrated in complete growth media for 1 h prior to seeding cells at a density of 1 x 10^5^ cells per scaffold. Cells were cultured for 2 d prior to fixing and staining for analysis. For experiments with cells seeded within fibers as a support, fiber solutions (10% v/v in PBS) were sterilized under germicidal light for 2 h, packed via centrifugation, and the PBS supernatant was exchanged with sterile C2C12 growth media for at least 2 h prior to packing for cell experiments. PHM-100 was prepared as described above and C2C12s were gently mixed into PHMs at a density of 1 x 10^7^ cells/ml and cultured for 1 d in a PDMS mold prior to fixing and staining.

### Cell staining

Prior to cell staining, C2C12s seeded on printed filaments were fixed in 10% v/v neutral buffered formalin for 15 min, permeabilized in 0.1% v/v Triton X-100 for 10 min, then blocked with 3% w/v bovine serum albumin for 2 h at room temperature. Cells were incubated with AlexaFluor-488 phalloidin for 1 h to visualize F-actin (1:600, Invitrogen). Samples were washed thrice in PBS to remove unbound molecules.

A similar protocol was utilized for experiments with C2C12s in PHMs supports. Cells were fixed for 1 h, permeabilized for 1 h, and blocked for 2 h using the same solution concentrations prior to tagging F-actin with AlexaFluor-488 phalloidin (1:200) for 2 h. Again, samples were washed in PBS thrice to remove unbound fluorophore.

### Microscopy and image analysis

Fluorescence microscopy for fiber segmentation and printed filament analysis was conducted on a Leica DMi8 widefield fluorescent microscope. For cell imaging on filaments or within supports, a Leica Stellaris Confocal microscope was utilized to image Z-stacks, with representative maximum projections shown here. Fiber length, diameter, and directionality, as well as cell orientation and area were all quantified using built-in ImageJ functionalities.

