## Supplemental Information for "Packed hydrogel microfibers as a granular medium"

### Table of Contents

**Figure S1. Methacrylate-functionalized hyaluronic acid (MeHA) $^1$H NMR spectrum**
**Figure S2. Quantification of fiber length and diameter**
**Figure S3. Rheological characterization of PHM-90 and PHM-80**
**Figure S4. UV crosslinking of PHM-100 High and PHM-100 Low**
**Figure S5. Modeling stress relaxation with a viscoelastic standard linear solid model**
**Figure S6. Fiber alignment during extrusion**
**Video S7. Extrusion manipulation**
**Video S8. Extrusion across table**

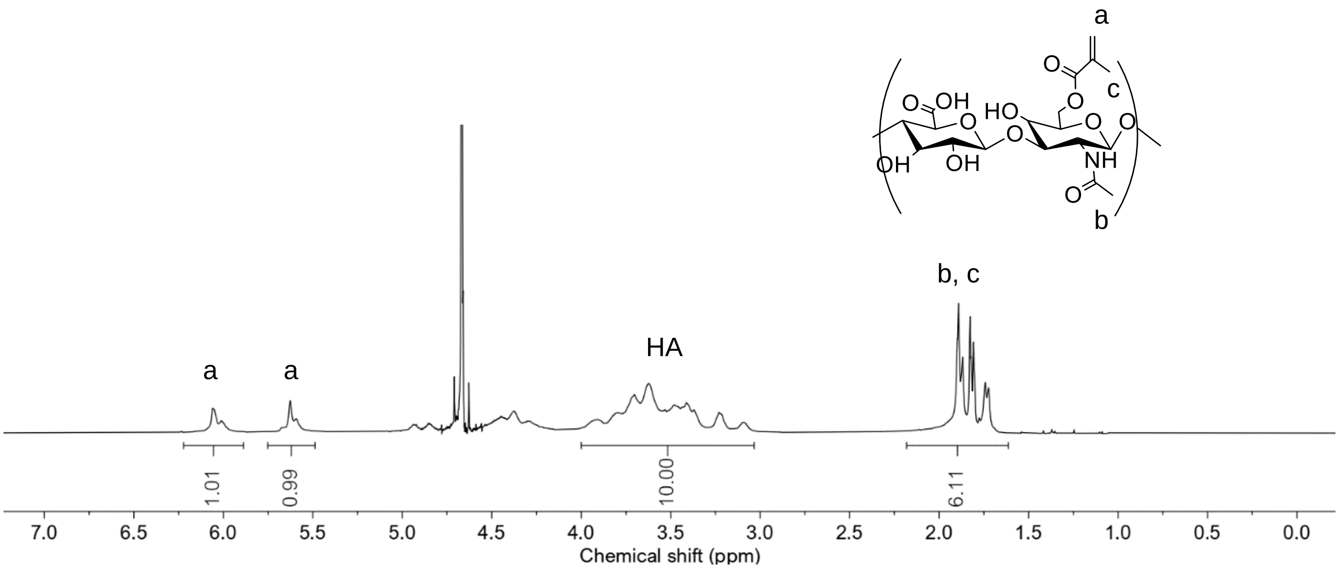


**Figure S1. Methacrylate-functionalized hyaluronic acid (MeHA) $^1$H NMR spectrum.** The resultant NMR spectrum is normalized to the 10 hydrogens on the standard HA backbone (~3.0-4.0 ppm), and the degree of modification is determined based on the integral of the peaks corresponding to the methacrylate groups (~5.5-6.25 ppm). The above spectrum illustrates a modification of ~100%.

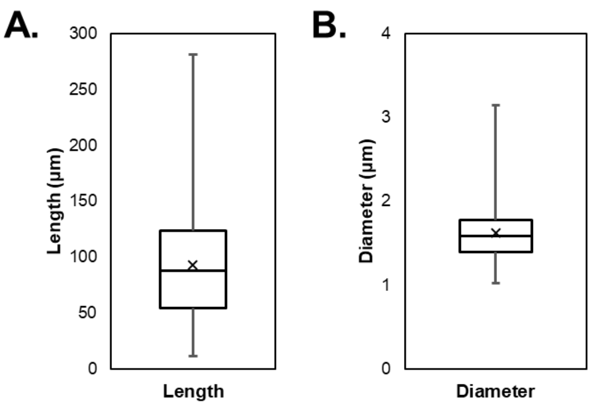


**Figure S2. Quantification of fiber length and diameter.** (A) Segmented fiber length was determined to be 93 $\pm$ 51 $\mu$m. (B) Segmented fiber diameter was determined to be 1.6 $\pm$ 0.3 $\mu$m. Fluorescent images of fiber solutions were utilized to quantify fiber segments.

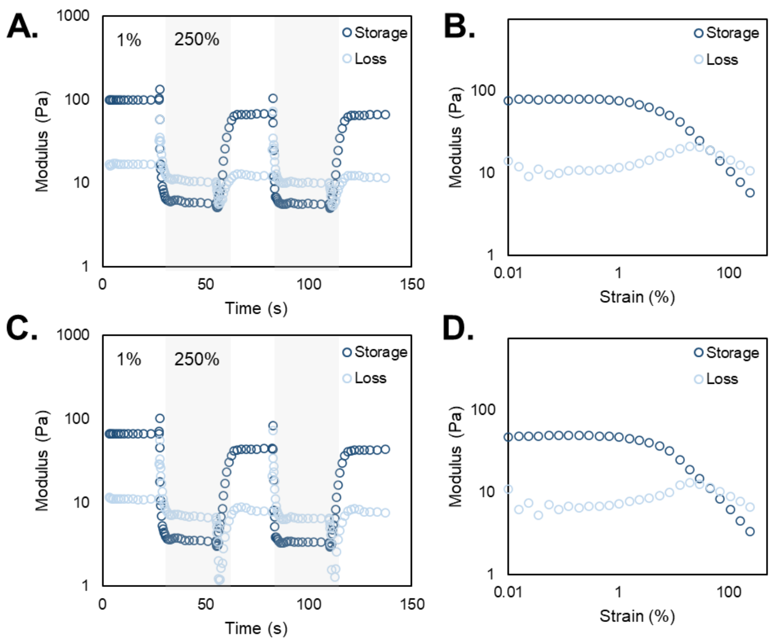


**Figure S3. Rheological characterization of PHM-90 (A-B) and PHM-80 (C-D).** (A, C) Shear-thinning and self-healing properties of PHM-90 and PHM-80, respectively. While modulus values are decreased compared to PHM-100, the trends are starkly similar, regardless of dilution. (B, D) Amplitude sweeps illustrate strain yielding properties of these diluted PHMs, with yield-strains significantly decreased compared to PHM-100 (shown in Figure 2, main text).

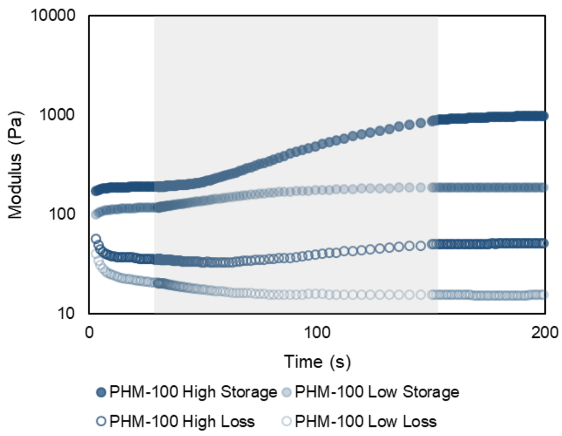


**Figure S4. UV crosslinking of PHM-100 High and PHM-100 Low.** Representative time sweeps including a UV cure step (shaded region). PHM-100 High, which has the most methacrylate groups available for crosslinking, exhibits a noticeable increase in storage modulus compared to the original pre-crosslinked state and the crosslinked PHM-100 Low as well. Importantly, both groups demonstrate a larger difference between their respective storage and loss moduli following the secondary crosslinking, which yields G’ values that are ~10x the G” values.

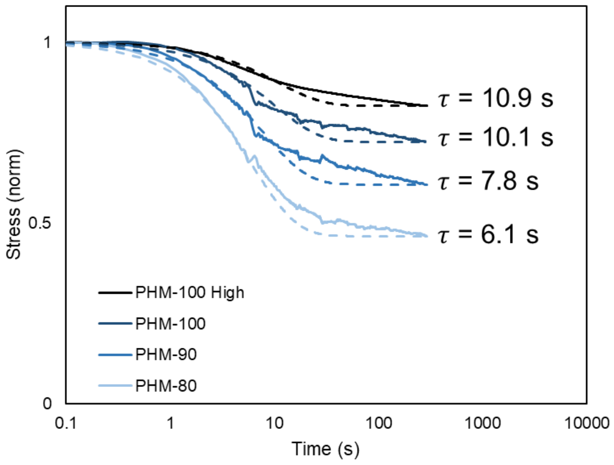


**Figure S5. Modeling stress relaxation with a viscoelastic standard linear solid (SLS) model.** Stress relaxation profiles of PHM-100, PHM-90, PHM-80, and PHM-100 High (solid lines) with corresponding SLS models (dashed lines). Representative relaxation time constants ($\tau$) are shown adjacent to each group.

The stress relaxation profiles of PHMs exhibit behaviors similar to viscoelastic solids. Interestingly, work by Wingert and coworkers[1] demonstrated that Nylon-11 nanofiber meshes dissipated stress similarly to PHMs, albeit at much longer time scales (on the order of minutes). Therefore, we leverage the same viscoelastic standard linear solid (SLS) model to analyze the stress relaxation of PHMs that was utilized to model the Nylon-11 fibers. The basic SLS model is shown below as Equation 1, and the resultant $\tau$ values are shown in the figure corresponding to the PHM group.

$$\frac{\sigma(t)}{\sigma_0} = \beta + (1-\beta)\, exp\left(-\frac{t}{\tau}\right) \quad (1)$$

Where:

$\tau$ = relaxation time constant

$$\beta = \lim_{t\to\infty} \frac{\sigma(t)}{\sigma_0}$$

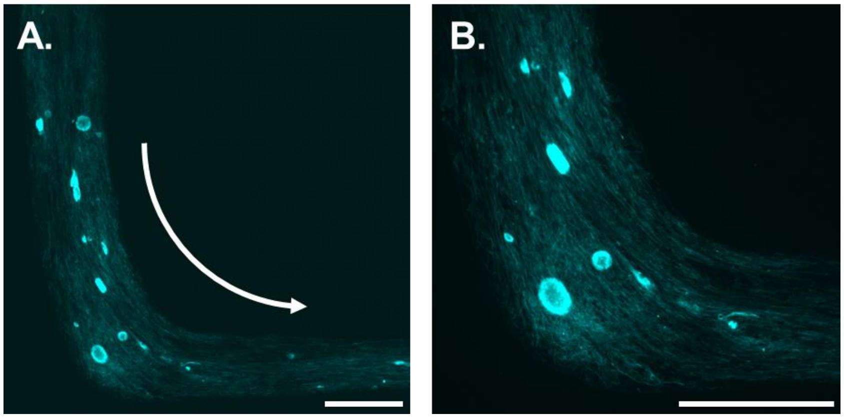


**Figure S6. Fiber alignment during extrusion.** The individual fiber segments within the PHM ink experience shear-induced alignment during extrusion printing processes. This microscale topography was demonstrated to provide contact guidance to C2C12 cells in the main text (Figure 6). This fiber alignment continues when the direction of extrusion changes. Shown in (A) and expanded in (B), fiber alignment continues with the curve as the direction of printing follows the white arrow in (A). It is important to note that the depicted filaments contain some aggregates of fibers. Overall, this demonstrated control over microscale topography enables the ability to arbitrarily define the surface topography of a substrate purely by extrusion design. *Scalebars = 500 µm.*

**Video S7. Extrusion manipulation.** Video corresponding to the images shown in Figure 5A of the main text. Video playback is 5x original speed.

**Video S8. Extrusion across table.** Video corresponding to the image of extruded filament across four corners of a table in Figure 5B of the main text. Video playback is 5x original speed.

**References:**

1 M. C. Wingert, Z. Jiang, R. Chen and S. Cai, *J. Appl. Phys.*, , DOI:10.1063/1.4973486.
